# Spikes meet Spins: Quantum-Native Neural Decoding for Ultra-Low-Latency Brain–Computer Interfaces

**DOI:** 10.64898/2026.04.09.717346

**Authors:** Gen Li, Yifei Ye, Haoyang Su, Ye Tian, Luyue Jiang, Yingkang Yang, Yaxuan Huang, Qi Gao, Kai Wen, Liuyang Sun

**Affiliations:** State Key Laboratory of Transducer Technology, Shanghai Institute of Microsystem and Information Technology, Chinese Academy of Sciences; Shanghai, 200050, China; Shanghai Innovation Institute; Shanghai, 200231, China; Beijing QBoson Quantum Technology Co., Ltd.; Beijing, 100015, China; School of Graduate Study, University of Chinese Academy of Sciences; Beijing, 100049, China

## Abstract

Brain – computer interfaces (BCIs) require rapid and accurate decoding of neural activity, yet conventional computing architectures face growing latency as neural recording scales. We demonstrate a quantum computing – enabled neural decoding approach using a physical 1000-qubit coherent photonic Ising machine, in which inference is performed through hardware energy relaxation rather than numerical computation. By mapping sparse neural spike patterns onto Ising Hamiltonians, our hardware-native Quantum Semi-Restricted Boltzmann Machine achieves up to 96.2% accuracy across public in vivo datasets spanning multiple species and modalities. We report hardware-verified median latencies of 0.075 ms— a tenfold speedup over GPUs—with complexity-invariant scaling. These results establish quantum computing as a viable pathway toward ultra-low-latency neural decoding for future BCI systems.

## Introduction

Rapid and accurate decoding of neural signals is key to BCIs, yet an escalating “latency crisis” threatens the scalability of the field^1–4^. While high-density electrode arrays now can record from thousands of neurons^5–10^, and deep learning architectures—such as Transformers— have reached high precision^1,4,11,12^, these advancements come at an unsustainable computational cost. As neural channel counts expand, the inference latency and energy consumption of conventional von Neumann hardware grow prohibitively^13^, even when accelerated by high-end graphics processing units (GPUs). This emerging accuracy-latency bottleneck prevents digital systems from keeping pace with the millisecond-scale dynamics of biological circuits^14,15^, imposing a fundamental constraint on the development of truly seamless closed-loop BCIs.

Quantum computing architectures offer a fundamentally different computational paradigm that remains largely unexplored in the context of neural engineering^16–19^. Specifically, Ising machines perform optimization by allowing a physical system to evolve naturally toward its ground state—transforming combinatorial inference into a process of energy relaxation rather than a sequence of digital operations^20–23^. Because neural spike patterns are inherently sparse, binary, and high-dimensional, they can be naturally mapped onto the spin-based representations used in Ising Hamiltonians and energy-based models such as Boltzmann Machines(BMs)^16,24–26^. Despite this intrinsic synergy, the prospect of a “quantum advantage” in neuroscience has remained a distant vision. To date, there has been no hardware-validated demonstration that the physics of a quantum system can be harnessed to decode complex, in vivo biological signals with the accuracy and speed required to challenge state-of-the-art (SOTA) conventional decoders.

Here, we bridge this gap by demonstrating a multi-modal, cross-species neural decoding framework executed on a 1,000-qubit coherent photonic Ising machine (CIM), transforming numerical inference into the physical energy relaxation of a photonic network. By encoding neural spike patterns into an Ising Hamiltonian, we utilize the coupled dynamics of optical parametric oscillators to evolve the system toward its lowest-energy configuration, thereby decoding neural intent via a hardware-native Quantum Semi-Restricted Boltzmann Machine (QSRBM). We validate this approach on diverse, multi-modal, cross-species, public available in vivo datasets, including visual stimulus–evoked activity in the mouse visual cortex and motor-intent population dynamics in the macaque motor and somatosensory cortices^27–30^. This broad validation demonstrates the generalizability of our “spikes-to-spins” interface across different neural coding regimes and species without data selection bias.

The QSRBM achieves decoding accuracy up to 96.2%, matching or surpassing state-of-the-art classical deep learning baselines. Crucially, within the available qubit budget, inference latency remains approximately constant with increasing neuron count, reaching a median of 0.075 ms (minimum 0.02 ms)—an order-of-magnitude speedup over GPU implementations. By uniting quantum computing and brain–computer interfaces—two of the most frontier technologies in modern science—this work establishes a shared physical framework for neural decoding, opening new avenues for both scalable BCI systems and application-driven quantum computing.

## Results

### Spikes-to-spins workflow for neural decoding on a photonic Ising machine

Decoding large-scale neural population activity is fundamentally constrained by the latency and power overhead associated with sequential digital computation. To overcome this limitation, we developed a quantum-native neural decoding framework where inference is performed through energy relaxation on a 1000-qubit CIM, rather than through iterative numerical optimization (Fig. 1).

**Fig. 1.**
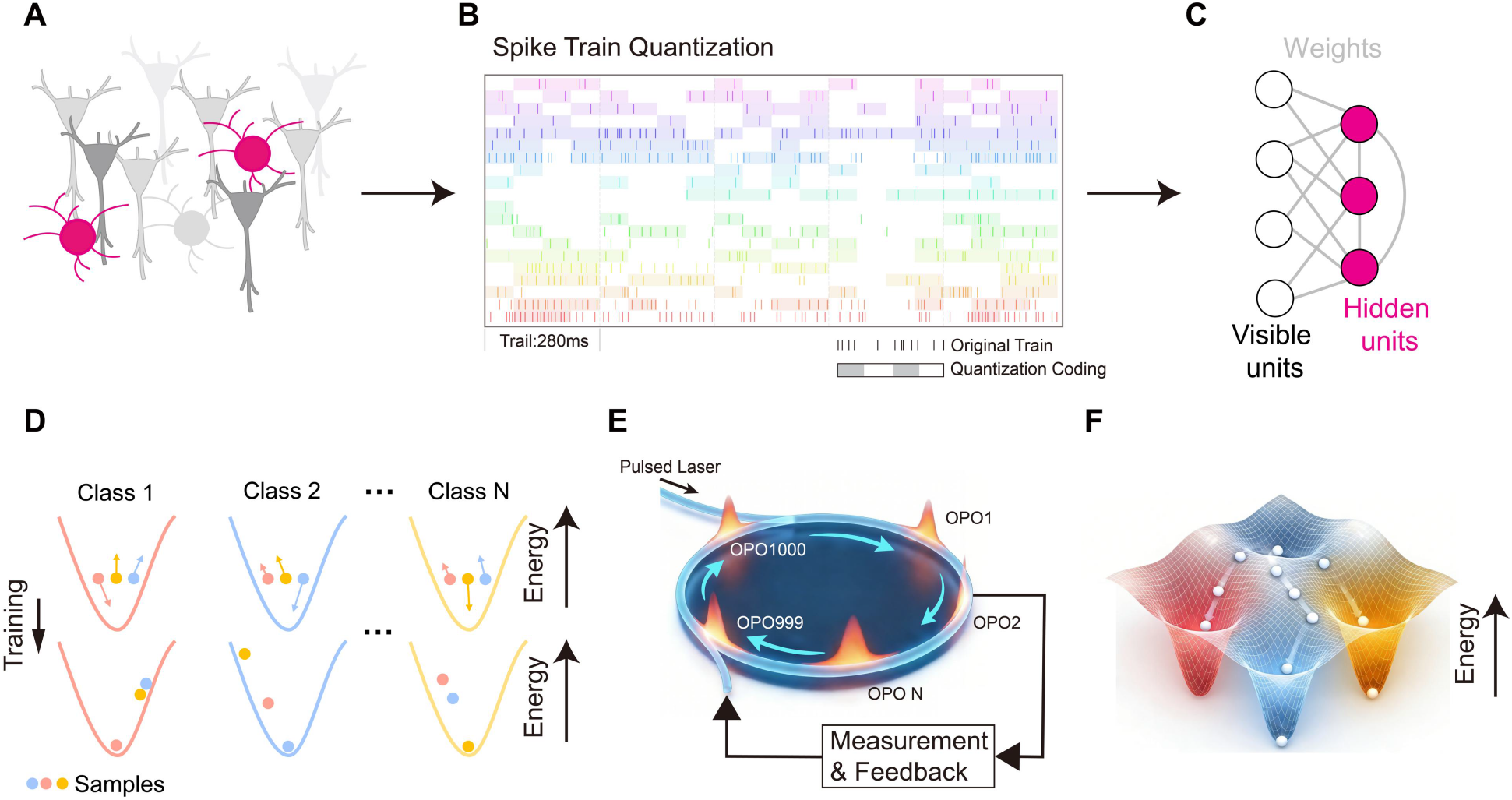
Spikes-to-spins workflow for neural decoding via physical energy relaxation. **(A, B)** Schematic of the “spikes-to-spins” interface, where multi-channel neural spike trains are mapped onto 4-bit discrete states compatible with the CIM. **(C)** Direct embedding of a QSRBM into a physical Ising Hamiltonian defined by programmable couplings ***J***_***ij***_ and local fields ***h***_***i***_. **(D)** Visualization of the learned energy landscape, showing the establishment of a pronounced energy gap (***ΔE***) and the transformation of the neural state space into attractor basins or “energy funnels”. **(E)** Conceptual diagram of the 1,000-qubit CIM hardware. **(F)** The autonomous relaxation process toward the global energy minimum (target class) stabilized by an optoelectronic feedback loop.

In this framework, simultaneously recorded neural spike trains are first mapped onto a low-dimensional discrete representation compatible with the spin-based architecture of the CIM (Fig. 1A, B)^20–22^. Given the finite quantum resources of the hardware, we implemented a Semi-Restricted Boltzmann Machine (SRBM) architecture^31^ in which qubits are divided between visible units—encoding neural inputs—and hidden units that capture latent cross-channel correlations. For a typical population of ∼200 neurons, we assigned a 4-bit discrete state to each neuron to preserve temporal information within the decoding window (Fig. 1B).

The core of our approach is the direct embedding of the SRBM into a physical Ising Hamiltonian, defined by programmable coupling terms Jij and local fields *h*i^20,22^. During training, class-labeled neural data are used to optimize these physical parameters, effectively sculpting the energy landscape of the photonic system (Fig. 1C). A critical outcome of this training is the establishment of a pronounced energy gap (ΔE) between the correct class and all competing alternatives (Fig. 1D). Notably, this process transforms the high-dimensional neural state space into a series of “energy funnels” or attractor basins. For a given neural input, the global energy minimum corresponds to its true class, while incorrect classes remain separated by an extensive energetic barrier (Fig. 1D bottom).

Following training, inference proceeds through the autonomous relaxation of the photonic network—stabilized by an optoelectronic measurement-and-feedback loop (Fig. 1E)—toward the lowest-energy configuration. When a neural sample is “dropped” into this physical energy landscape, the robust energy gap ensures that the system settles into the correct target basin, while random noise and competing patterns remain trapped at higher energies (Supplementary Fig. S1). Final classification is determined by identifying the global minimum across the class-specific Hamiltonians.

Beyond discrete classification, this physical relaxation generates a structured latent representation of neural activity. As a representative example, we visualize the energy-vector manifold for a 16-class decoding task, where test samples corresponding to distinct stimuli form well-separated clusters in a reduced-dimensional space (Supplementary Fig. S2). This organization emerges directly from the learned energy landscape, without the need for explicit feature engineering or iterative optimization.

Together, this workflow establishes a direct mapping from biological spike dynamics to physical energy minimization, allowing the underlying physics of the light field to perform inference without the overhead of algorithmic control flow.

### Decoding accuracy across sensory and motor cortical datasets

To assess the functional reliability and generality of this quantum-native decoding paradigm, we benchmarked its performance across multiple species, cortical areas, and task modalities, and compared it against state-of-the-art classical neural decoders executed on GPU hardware.

#### Multi-modal neural decoding across species and tasks

A critical requirement for this hardware-integrated pipeline is the efficient “quantization” of biological spike trains into discrete spin configurations compatible with the CIM architecture. To identify the optimal interface, we systematically compared six biologically motivated quantization strategies: (1) 4-segment Temporal, (2) Log-Temporal, (3) Truncated, (4) Onset + Sustained, (5) Latency–Rate, and (6) Rate–Slope(Fig. 2A, B).

**Fig. 2.**
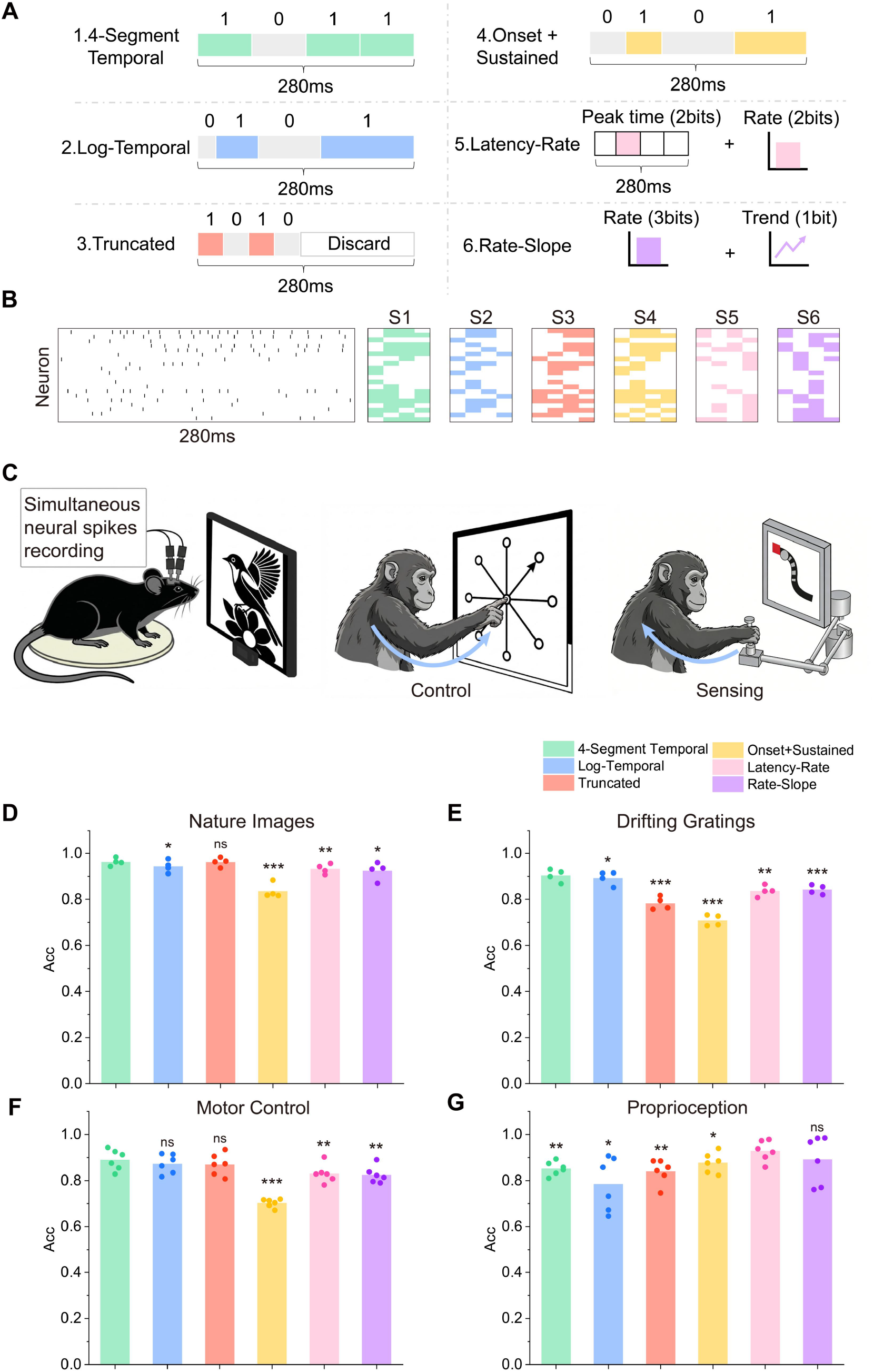
Multi-modal spike quantization and task validation. **(A, B)** Comparison of six biologically motivated 4-bit quantization strategies (S1–S6) used to unify biological spikes into quantum-hardware-compatible spin configurations. **(C)** Overview of cross-species and multi-modal validation using publicly available in vivo datasets, including mouse visual cortex (V1) and macaque motor and somatosensory cortices. **(D–G)** Decoding accuracy across four representative tasks. For mouse tasks (D and E, natural images and drifting gratings), data were collected from n=4 mice. Individual dots represent the mean decoding accuracy for each recording session, and bars indicate the average of these session means. For monkey tasks, two independent datasets were used: motor control (F) was tested in n=2 rhesus macaques, and proprioception (G) was tested in n=2 cynomolgus macaques. Due to the limited sample size per dataset, three independent trials with different random seeds were conducted for each condition; dots represent the mean accuracy per trial. Statistical significance was assessed using a paired t-test to compare the performance of different quantization strategies. Significance levels are indicated as *P < 0.05; **P < 0.01; ***P < 0.001; ns, not significant.

We first evaluated the robustness of our quantum-native framework using diverse, publicly available in vivo recordings. By utilizing open-access datasets, we ensure that our results are cross-validated on standardized neural benchmarks, precluding concerns regarding data selection bias and enabling direct comparison with state-of-the-art classical models. These datasets encompass multiple neural modalities, including visual stimulus-evoked activity from the mouse primary visual cortex (V1) and motor-intent and somatosensory population dynamics from rhesus macaques and cynomolgus macaques(Fig. 2C).

Across all species and tasks, quantization strategies that preserved fine-grained temporal structure consistently outperformed simplified rate-based representations. In the mouse visual cortex, the 4-segment Temporal and Log-Temporal encodings achieved peak decoding accuracies of **96.2%** for natural images and **90.4%** for drifting gratings(Fig. 2D, E). Notably, these temporal strategies maintained a significant performance edge even as the stimulus complexity shifted from static images to dynamic gratings.

To test the clinical relevance of our framework for BCIs, we extended our validation to macaque population dynamics during a reaching task. For motor control decoding (identifying reach direction), the 4-segment Temporal and Truncated encodings reached accuracies of **∼89%**(Fig. 2F). Interestingly, in the decoding of proprioceptive feedback—a task typically characterized by high-frequency rate modulations—the Latency–Rate strategy emerged as top performers, achieving **∼92.9%** accuracy(Fig. 2G). We hypothesize that this divergence reflects the distinct coding mechanisms of mechanosensory pathways: unlike cortical processing which relies heavily on precise population spike-timing, proprioceptive feedback optimally encodes continuous kinematic variables and rapid mechanical transitions through a combination of robust sustained rates and precise initial latencies.

#### Performance comparison with state-of-the-art classical decoders

To benchmark the efficacy of the quantum-native decoder, we compared the QSRBM against a suite of state-of-the-art classical architectures, including Convolutional Neural Networks (CNN), Long Short-Term Memory (LSTM) networks, Transformers, and Conformers^32^. To ensure a rigorous comparison, all classical baselines were executed on high-end GPU hardware and trained using the same unified 4-bit spike quantization, ensuring that any performance gains were attributable to the physical inference mechanism rather than preprocessing advantages.

The QSRBM consistently outperformed all classical models across both species and every recording modality tested. In the mouse visual cortex tasks, the QSRBM achieved a peak accuracy of **96.2%** for natural images, surpassing the Conformer (**91.1%**) and significantly outperforming the Transformer (**86.9%**) and CNN (**81.8%**) baselines(Fig. 3A). For the dynamic drifting grating task, the QSRBM maintained a high accuracy of **90.4%**, representing a substantial improvement over the Transformer (**81.1%**) and nearly doubling the performance of the LSTM (**60.9%**)(Fig. 3B).

**Fig. 3.**
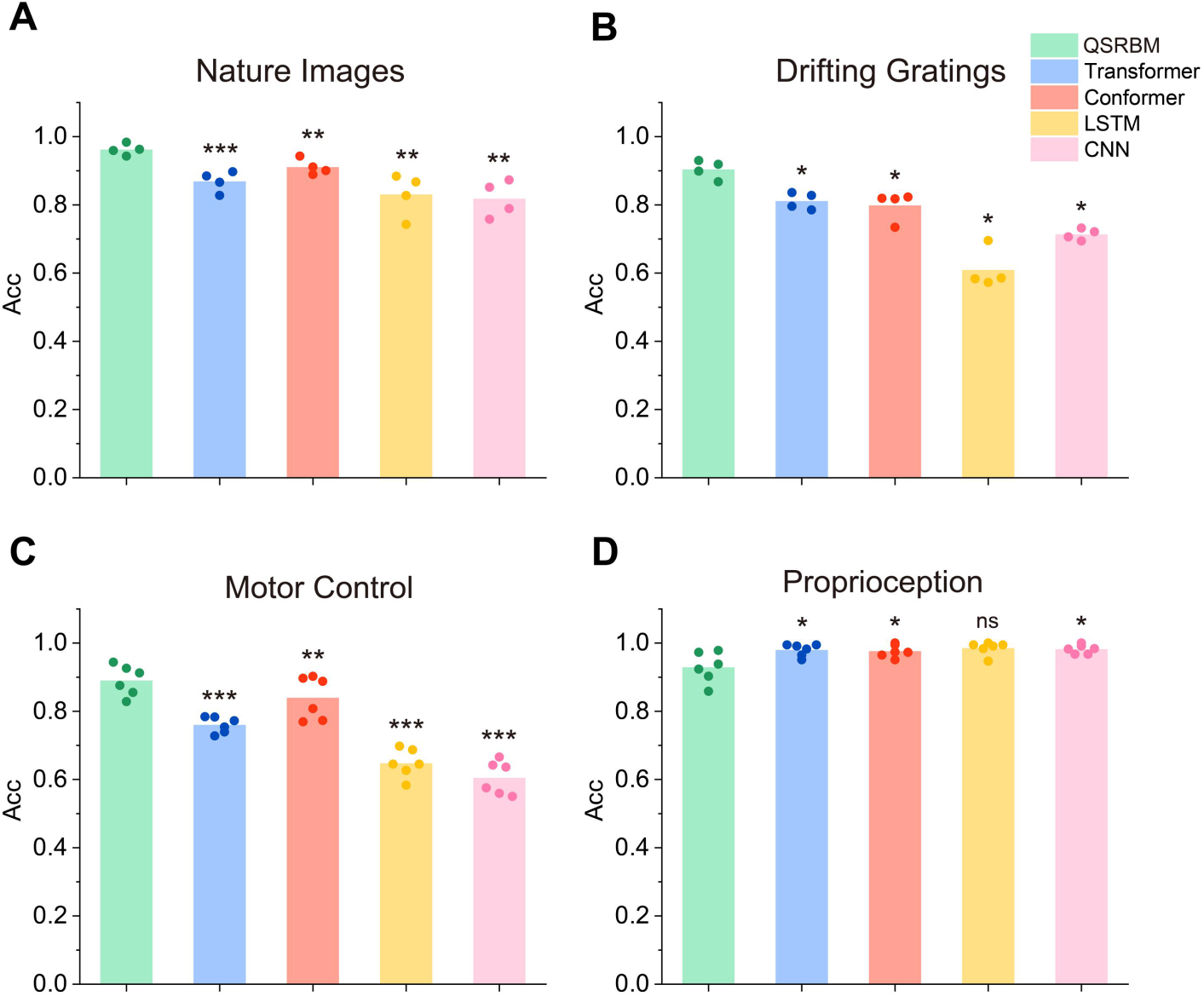
Performance comparison against state-of-the-art classical decoders. **(A, B)** Head-to-head decoding accuracy comparison of the quantum-native QSRBM against GPU-accelerated classical architectures (Transformer, Conformer, LSTM, and CNN) across four benchmark tasks. For mouse visual stimuli (A, natural images and B, drifting gratings), data were collected from n=4 mice; individual dots represent the mean decoding accuracy for each recording session, and bars indicate the average of these session means. For monkey tasks, independent datasets were used: **(C)**, motor control in n=2 rhesus macaques, and **(D)**, proprioception in n=2 cynomolgus macaques. Due to limited sample sizes, three independent trials with different random seeds were conducted for monkey tasks; dots represent the mean accuracy per trial. Statistical significance was assessed using a paired t-test (***p < 0.001, **p < 0.01, *p < 0.05, ns = not significant), confirming that performance gains arise from the physics-native inference mechanism.

Detailed analysis of the classification errors via confusion matrices reveals that the QSRBM captures the underlying semantic structure of the stimuli (Supplementary Fig. S3). For instance, in the drifting grating task, residual errors primarily occur between oppositely moving directions, reflecting intrinsic ambiguities in neural population responses rather than a failure of the quantum algorithm.

Extending the evaluation to the motor control task, the QSRBM continued to demonstrate robust decoding capabilities, achieving an accuracy of **89.0%**. This performance notably surpassed the Conformer (**83.94%**) and the Transformer (**75.9%**), while the LSTM and CNN baselines exhibited significantly lower accuracies, further validating the quantum-native decoder’s superiority in handling complex motor cortex dynamics(Fig. 3C). Confusion matrices for the motor task show that misclassifications are largely confined to adjacent reach trajectories (Supplementary Fig. S4), suggesting that the learned energy landscape of the CIM effectively preserves the directional manifold of the motor cortex.

However, the proprioception task revealed a distinct performance shift. While the QSRBM maintained a strong accuracy of **92.9%**, the classical models exhibited a marked surge in performance, achieving an average accuracy of **98.1%**(Fig. 3D). We attribute this inversion to the drastic reduction in neuronal population size. Unlike the visual cortex tasks (∼200 neurons) or motor control (194 neurons), the proprioception recordings averaged only ∼50 neurons. In this low-dimensional regime, the fixed architectural complexity of the classical models became highly efficient, allowing their extensive parameter spaces to easily model the simplified input distribution. This contrast highlights a critical insight: while the QSRBM excels in extracting patterns from high-dimensional, complex neural representations where classical models struggle, the massive parameter capacity of classical deep learning models can offer marginal advantages in data-scarce, low-dimensional scenarios, effectively saturating the decoding ceiling.

#### Robustness of quantum inference under input quantization

While the QSRBM demonstrates superior performance on unified inputs, a central question remains regarding the impact of the data representation itself. Integrating quantum hardware with biological systems necessitates a transition to discrete, hardware-compatible formats. To isolate the impact of our 4-bit “spikes-to-spins” quantization, we performed an additional control experiment comparing the QSRBM (operating on 4-bit data) against state-of-the-art classical models operating on the original, uncompressed spike trains.(Supplementary Fig. S5)

Under these unconstrained conditions, classical models achieved accuracies comparable to the QSRBM. However, this result highlights a fundamental efficiency trade-off: while classical architectures can recover high performance when given access to high-dimensional, raw temporal features, the QSRBM maintains peak decoding accuracy under extreme input compression. Such efficiency is critical for power-constrained BCIs, where the computational overhead of processing raw, high-dimensional spike streams is often prohibitive.

We observed a consistent performance hierarchy across all datasets: the QSRBM operating on unified 4-bit data achieved accuracies comparable to GPU-based models trained on raw, un-unified data—with only a marginal 2% difference observed in the dynamic grating task—while GPU-based models trained on the same 4-bit data exhibited the lowest performance. This hierarchy indicates that the quantization process itself is not responsible for the QSRBM’s success. Instead, the physical energy-minimization dynamics of the Ising machine enable the effective extraction of task-relevant neural structures even under reduced input precision.

These results underscore the robustness and expressivity of physics-native inference. The QSRBM achieves state-of-the-art accuracy using a fixed, low-bit representation that is directly compatible with ultra-fast hardware, providing a pathway for high-performance decoding without the need for complex, power-hungry preprocessing pipelines.

### Hardware-validated inference latency

To determine if quantum-native inference provides a fundamental advantage over classical architectures, we directly measured the end-to-end inference latency and energy consumption of the QSRBM compared to GPU-accelerated state-of-the-art models.

#### Mechanistic basis of ultra-low latency

Unlike the iterative numerical optimization required by digital processors, inference on the photonic Ising machine is a physical process of energy relaxation. The hardware utilizes a 1km fiber-optic ring in which optical pulses circulate; each “iteration” in the Hamiltonian evolution corresponds to a single round-trip through this fiber length. We monitored the normalized Hamiltonian during this process and observed a rapid, monotonic decay toward the ground state. The system typically settles into a stable minimum in tens of iterations; evaluation across 36,333 quantized neural samples demonstrated a median “number of shots” of 15.0 and a mean of 26.2 (Fig. 4A, B). This confirms that the physical convergence is both extremely rapid and highly reliable.

**Fig. 4.**
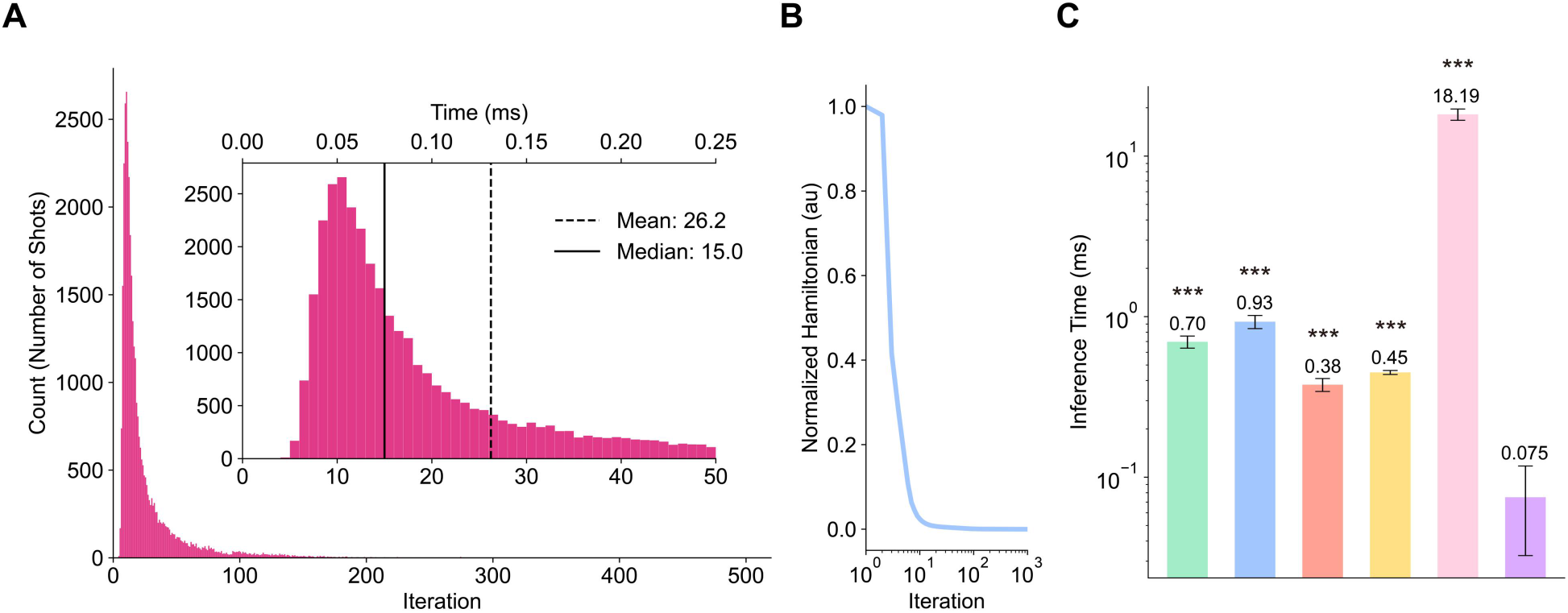
Hardware-validated latency, scaling, and energy efficiency. **(A)** Distribution of the number of “shots” (iterations) required to reach the ground state during physical relaxation, derived from 36,333 hardware decoding trials; inset shows the mapping of iterations to hardware time (median = 15.0 iterations, mean = 26.2 iterations). **(B)** Monotonic decay of the normalized Hamiltonian during the 1km fiber loop evolution, illustrating the rapid convergence toward the target class basin (data derived from the same 36,333 trials). **(C)** Measured end-to-end inference latency, highlighting the QSRBM’s 0.075 ms average speed — an order-of-magnitude improvement over GPU-accelerated decoders. Measurements were based on n=36,333 trials for QSRBM and n=30,000 trials for GPU models. Statistical significance was assessed using an unpaired t-test (Welch’s t-test), confirming significant superiority (***p < 0.001).

#### Hardware-native speed and constant-time scaling

This physical efficiency translates directly into superior decoding speeds. To ensure a rigorous comparison, inference latencies were measured as the pure processing time for each individual sample—utilizing CUDA.event for GPU baselines to capture kernel execution exclusively, thereby excluding any preprocessing overhead; an equivalent strict methodology was applied to the QSRBM Inference latency on the GPU increased significantly with both model complexity and neural population size. For a population of ∼200 neurons, classical models such as Transformers and Conformers required 0.70 ms and 0.93 ms per inference, respectively. To isolate the source of our speedup, we benchmarked a digital SRBM on the same GPU hardware. Notably, the digital SRBM exhibited the highest latency of all models tested (∼18 ms), confirming that the SRBM architecture itself is prohibitively expensive when executed on von Neumann hardware (Fig. 4C).

In contrast, the same SRBM architecture executed on the photonic Ising machine achieved an average latency of 0.075 ms, with minimums as low as 0.02 ms. This represents an approximately 240-fold speedup over its digital SRBM counterpart and an order-of-magnitude improvement over the fastest GPU-accelerated SOTA models. These results demonstrate that the observed gains in efficiency are not properties of the SRBM model itself, but are strictly derived from the physics of the quantum-native CIM hardware.

Furthermore, while GPU-based latency grows with input dimensionality, the QSRBM exhibits complexity-invariant scaling. Because all 1000 qubits evolve in parallel within the photonic network, latency remains approximately constant as the neural population size increases, provided the task remains within the hardware’s qubit budget.

## Discussion

In this work, we demonstrate that neural decoding can be executed as a physical process on quantum hardware, rather than as a sequence of numerical operations on electronic processors. By mapping sparse neural spike patterns onto an Ising energy landscape, we utilize the intrinsic relaxation dynamics of a 1000-qubit coherent photonic Ising machine to perform inference.

A defining distinction of this framework is its departure from the hybrid quantum-classical pipelines that have characterized early explorations in this field^33,34^. Previous attempts to integrate quantum hardware into neural decoding have largely relied on hybrid models, where conventional deep learning architectures perform the bulk of the computation, with a quantum layer appended only for final-stage refinement. Furthermore, many of these efforts have remained confined to theoretical simulations rather than physical execution.

In contrast, our approach represents a quantum-native architecture; both training and inference are executed directly on the physical Ising hardware. By eliminating the need for conventional deep learning algorithms entirely, we bypass the digital preprocessing bottlenecks that typically negate the speed advantages of quantum hardware, allowing the intrinsic parallelism of the photonic system to determine the decoding outcome. This hardware-validated implementation of a QSRBM using in vivo neural data allows the system to reach decoding accuracies of up to 96.2% across diverse datasets—matching or surpassing state-of-the-art models like Transformers and Conformers.

The transition from algorithmic simulation to physical computation provides a transformative advantage in speed and efficiency. We measured a hardware-verified average latency of 0.075 ms, representing a 240-fold speedup over its digital SRBM counterpart and a tenfold improvement over GPU-accelerated classical models. Crucially, the CIM exhibits complexity-invariant scaling; because the system evolves in parallel, inference latency remains approximately constant even as neural population sizes increase.

The requirement to discretize neural activity into hardware-compatible 4-bit representations introduces an important trade-off. However, by benchmarking against classical models operating on raw spike data, we show that the energy-based dynamics of the Ising machine effectively extract task-relevant structures even under extreme data compression. This high-fidelity performance with quantized inputs highlights the robustness of physical inference for sparse biological signals.

While this study emphasizes the latency advantages of quantum-native decoding, several limitations warrant further consideration. Our evaluation utilized offline datasets and a fixed 1000-qubit budget. Future work will explore online decoding, adaptive temporal encoding, and integration with larger-scale quantum hardware. Additionally, while this study emphasizes latency, a comprehensive system-level analysis of energy efficiency remains an important direction for further investigation.

More broadly, this work bridges two frontier fields—quantum computing and brain– computer interfaces—by demonstrating how physical quantum systems can directly process biological neural information. As neural interfaces continue to scale in channel count and complexity, approaches that allow physics itself to perform inference may provide a viable path beyond the limitations of conventional computing architectures, enabling faster, more efficient, and more naturalistic interactions between brains and machines.

## Methods

### Mouse Visual Cortex Dataset

This dataset^27^ utilized in vivo neural recordings from the primary visual cortex (V1) of four male mice (3–6 months old). Flexible electrode arrays were chronically implanted targeting the V1 visual areas. Neural signals were acquired at 30 kHz using an Intan RHD recording system, and spike sorting was performed using Mountainsort4^35^ with strict quality control criteria for single-unit identification. Putative single units were tracked across days based on waveform similarity and spatial location estimates. Visual stimuli were presented on a widescreen monitor placed 21 cm from the animal. The study employed two stimulus sets: 1.Drifting Gratings (DG): Full-contrast sinusoidal gratings drifting in 16 equally spaced directions (spatial frequency: 0.05 cycles per degree; temporal frequency: 2 Hz). Each stimulus was presented for 520 ms, followed by a 500 ms gray screen. A total of 800 trials were presented daily. 2.Natural Images (NI): A set of 100 grayscale images from the ImageNet database. Each image was presented for 245 ms with no inter-stimulus gray screen. For our analysis, we used the data corresponding to the first 16 images from this natural image stimulus set.

The order of stimulus conditions was randomized within each block. Stimulus onsets were precisely synchronized with neural recordings using a light-to-frequency converter attached to the screen corner. From this dataset, neural activity from four mice was included in the final analysis.

### Motor Control Dataset

Neural data^30^ were obtained from two male rhesus macaques (Monkeys Jenkins and Nitschke) performing a delayed center-out reaching task. In the task, monkeys initiated a reach from a central position to one of multiple peripheral targets after a variable delay period (400– 1000 ms). The dataset included recordings from primary motor cortex (M1) and dorsal premotor cortex (PMd), utilizing both conventional single-electrode techniques and chronically implanted 96-electrode arrays (Blackrock Microsystems). A total of 194 well-isolated single units were recorded from each monkey and used for subsequent analysis.

Data preprocessing was performed using custom Python scripts. Only successful trials (task_success=1) from the center-out task variant (maze_num_barriers=0) were selected. To ensure precise directional tuning, reach endpoints were categorized into eight standard directions (0 °, 45 °, 90 °, etc.) with a strict tolerance of ± 15 ° . Trials with ambiguous reach vectors (deviating >15 ° from any standard direction) or minimal movement amplitude (<5 mm) were excluded. One direction (Dir 6, 225°) was additionally removed due to insufficient trial count.

For each trial, spike trains were aligned to movement onset over a 600-ms window (100 ms pre-movement to 500 ms post-movement). Simultaneously, hand position trajectories were resampled via linear interpolation to align with the spike bins and were centered relative to the movement onset position. The dataset was partitioned for model training and validation, ensuring balanced representation across the seven included reach directions.

### Proprioception Dataset

Neural data^28^ were sourced from two male macaques (Monkeys C and H) performing a classic center-out reaching task using a planar manipulandum. The task involved holding a cursor at a central target and reaching to one of eight peripheral targets after a go cue. In half of the trials (passive trials), a 2 N perturbation was applied to the hand during the hold period; however, only active (non-perturbed) trials were used in this study.

Neural activity was recorded from a 96-electrode Utah array (Blackrock Microsystems) implanted in the arm representation of somatosensory area 2. Threshold crossings were subsequently sorted offline using Plexon Offline Sorter to isolate putative single units. To ensure data quality, sorted units with firing rates less than 0.1 Hz were excluded from analysis. Due to recording variability across days and stringent quality filtering, approximately 50 stable single units per monkey were retained for the final dataset.

Data preprocessing was conducted using custom Python scripts. For each trial, spike trains were aligned to the go cue (target onset) a 600-ms window (100 ms pre-movement to 500 ms post-movement). Simultaneously, hand position trajectories were resampled to align with the spike bins. Reach direction labels (0 – 7) were computed based on the angle of the vector connecting the start and end points of the trajectory, using the mean of the first and last five time bins to minimize noise.

To address imbalances in the distribution of directions within the dataset, a final directional filtering step was applied. Specifically, only trials corresponding to the four most well-represented directions for each monkey were retained, ensuring a balanced dataset for model training and evaluation.

### Quantization Strategies for Spike Data

To enable efficient mapping of neural spiking activity onto quantum hardware, we evaluated six distinct 4-bit quantization strategies that transform continuous spike trains into discrete spin configurations. These strategies were designed to capture different temporal and dynamic features of neural responses, balancing biological plausibility with computational efficiency. All strategies operated on spike count matrices within a defined temporal window (e.g., [-100 ms, 500 ms] relative to movement onset). For each strategy, a baseline statistic was first computed from the training set to ensure generalization.

#### 4-segment Temporal Strategy

This strategy partitioned the temporal window into four equal segments, capturing the coarse temporal evolution of firing rates. For each neuron, a baseline firing rate b was computed as the global mean across all training trials and time bins. Within each segment s, the mean firing rate μ_s_ was calculated. A binary digit was assigned per segment:

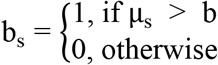

The four resulting bits were concatenated to form a 4-bit integer, encoding a simple temporal profile relative to the neuron’s average activity. This approach is inspired by the phasic-tonic response patterns observed in cortical neurons.

#### Log-Temporal Strategy

To reflect the non-linear temporal processing dynamics of the visual system, where early response components often carry more information, this strategy used logarithmically spaced time segments. The temporal window was divided into four segments with boundaries at 0 ms, 50 ms, 200 ms, 350 ms, and 520 ms post-stimulus as for motor control dataset. Within each segment, the same comparison against a global baseline b (as in Strategy 1) was performed to generate each bit. This encoding emphasizes fine temporal resolution during the initial response phase and coarser resolution later, aligning with the temporal dynamics of sensory processing.

#### Truncated Strategy

Focusing on the initial phase of the neural response, which is often most discriminative, this strategy considered only the first 260 ms of the temporal window as for motor control dataset. This window was divided into four equal segments, and within each, the mean firing rate was compared to a baseline b, calculated solely from the truncated portion of the training data. This approach reduces computational load and may be particularly relevant for decision-making processes that rely on early sensory evidence.

#### Onset + Sustained Strategy

This strategy explicitly separated the transient onset response from the sustained activity, a fundamental dichotomy in neural coding. The temporal window was split at 100 ms into an “onset” period (0-100 ms) and a “sustained” period (100-500 ms). Each period was further divided into two equal sub-periods. For each sub-period, the total spike count within it was compared to half of the total spike count for the entire period. For the onset period:

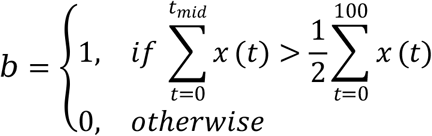

and similarly for the second sub-period and the sustained period. The four resulting bits provided a code reflecting the temporal distribution of spikes within the onset and sustained phases.

#### Latency–Rate Strategy

This strategy encoded two key features: response latency and firing rate amplitude. The temporal window was divided into four equal segments. Latency was encoded by the index k (0 to 3) of the segment with the highest total spike count:

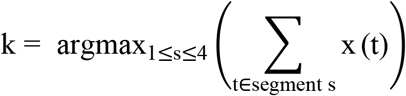

representing the timing of the peak response. Amplitude was encoded by normalizing the total spike count across the entire window for each neuron across trials:

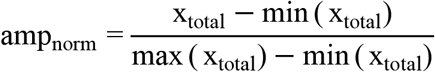

This normalized value was then discretized into two bits (0 to 3). The final 4-bit code was formed by combining the two latency bits and two amplitude bits. This captures both “when” and “how strongly” a neuron responds, which are critical features in sensory and motor coding.

#### Rate–Slope Strategy

This strategy encoded the overall firing strength and its temporal trend. Firing rate was quantified using three bits, derived from the same self-normalized amplitude as in Strategy 5, discretized into eight levels. The trend (slope) was encoded as a single bit by comparing the total spike count in the first half of the window to that in the second half:

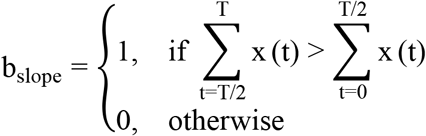

This bit distinguishes neurons with increasing versus decreasing firing rates over time. The combined 4-bit code thus captures both the overall excitability and the dynamic nature of the neural response. Trends in firing rate are indicative of adaptation, anticipation, or movement-related modulations.

### Quantum Computing Hardware

Decoding computations were performed using a CPQC-1000 coherent Ising machine (CIM) provided by Beijing QBoson Quantum Technology^22^. The CIM operates as a hybrid quantum-classical system at room temperature, utilizing an optical fiber loop to generate and store degenerate optical parametric oscillator (OPO) pulses as quantum bits (qubits). This architecture inherently supports fully connected Ising models. The computational process relies on a measurement-feedback loop: the phase and amplitude of the OPO pulses are measured by balanced homodyne detectors (BHD), and subsequently, feedback signals are calculated based on the target Ising Hamiltonian. These signals are applied via optical modulators to induce interference with the pulses in the loop, driving the system towards the ground state of the Hamiltonian. This physical mechanism enables the efficient minimization of the Ising energy function, providing solutions to the decoding optimization problem.

### Model Architecture and Ising Mapping

The decoding model was implemented as a Semi-Restricted Boltzmann Machine^31^ (SRBM) with a fully connected hidden layer, tailored for the quantum annealing architecture. The network topology comprises a visible layer v and a hidden layer ***h***.

#### Network Topology

The visible layer serves as the input interface for neural data. Given a population of ***M*** neurons where each neuron’s activity is quantized into ***b*** bits (e.g., ***b=4***), the visible layer consists of ***N***_***v***_ ***= M * b*** nodes. Each node is a binary unit ***vi∈{0, 1}***. The hidden layer serves as a feature extractor, with the number of hidden nodes ***N***_***h***_ set to approximately one-tenth of the visible nodes. Crucially, while the visible and hidden layers are fully connected via a weight matrix ***W***, the hidden layer itself is fully intra-connected via a coupling matrix ***J***. This structural modification transforms the model into a form compatible with the Ising Hamiltonian.

#### Energy Function

The state of the network is defined by the configuration ***(v, h)***. The energy function of the model is given by:

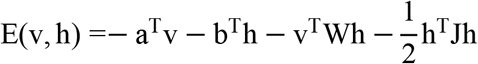

where ***a*** and ***b*** are the bias vectors for the visible and hidden layers, respectively; ***W*** represents the visible-hidden weights; and ***J*** is the symmetric coupling matrix for the hidden units.

#### Ising Model Mapping

To execute the decoding task on the CIM, the SRBM state was mapped to an Ising spin configuration. The binary states of the visible and hidden units ***(v, h)*** were collectively mapped to a spin vector ***σ∈{−1***,***+1}***^***N***^, where ***N = N***_***v***_ ***+ N***_***h***_. The mapping follows the transformation ***σ=2x−1*** for a binary unit ***x***. The optimization objective corresponds to finding the ground state of the following Ising Hamiltonian:

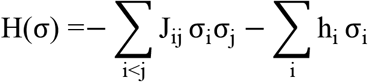

Here, the interaction matrix ***J***_***ij***_ encompasses the visible-hidden weights ***W*** and hidden-hidden couplings ***J***, while the local fields hi correspond to the biases ***a*** and ***b***. By minimizing this Hamiltonian via the CIM’s physical evolution, the system converges to a low-energy state that represents the optimal solution to the decoding problem.

### Model Training and Quantum Inference

The Quantum Semi-Restricted Boltzmann Machine (QSRBM) model was initially pre-trained on a classical graphics processing unit (GPU) to learn the data distribution efficiently. Given the complex intra-layer connections within the hidden layer, we employed a mean-field approximation algorithm with 20 iterations to estimate the posterior probabilities during training. Following the unsupervised pre-training, a discriminative fine-tuning phase was conducted using a low learning rate. This step aimed to sharpen the decision boundaries between different classes, thereby maximizing classification accuracy.

For inference on the CIM, the trained model parameters and the input test samples were transformed into Ising coupling matrices using the kaiwu Python library provided by QBoson Quantum Technology. Specifically, for a given test sample, Ising matrices corresponding to each candidate class were constructed and integrated into a unified Hamiltonian matrix. This matrix was then submitted to the CIM hardware via the kaiwu SDK. The CIM performed a physical evolution to search for the ground state, and the resulting low-energy configuration was mapped back to the predicted class label.

### Performance Evaluation Metrics

To rigorously compare the computational efficiency of the proposed QSRBM against classical GPU-accelerated decoders, we established a unified framework for measuring inference latency .

For GPU-based models, all experiments were conducted on an NVIDIA GeForce RTX 4090 GPU. The inference time per single sample was measured using ***CUDA*.*event***, which provide high-precision timing on the GPU device. The reported latency for each model represents the average wall-clock time over 30,000 individual inference trials. For the QSRBM, the CIM returns the number of iterative shots required for the system to converge to a stable ground state. Since the duration of each shot is a fixed physical constant of the hardware, the total inference time for each of the 36,333 samples was calculated by summing the total shot counts and multiplying by the fixed period per shot.

## Author contributions

Conceptualization: LS, GL

Methodology: GL, LS, LJ

Investigation: GL, LS, HS

Visualization: GL, LS, YY, YH

Funding acquisition: LS, KW, QG

Project administration: LS

Supervision: LS

Writing – original draft: LS, GL

Writing – review & editing: LS, GL, YT, YY, YH

## Data, code, and materials availability

All data needed to evaluate the conclusions in the paper are present in the paper and/or the supplementary materials. Additional data related to this paper and the customized scripts are available from the corresponding author upon reasonable request.

## Supplementary Materials

Figs. S1 to S5

## References

1. Silva, A. B., Littlejohn, K. T., Liu, J. R., Moses, D. A. & Chang, E. F. The speech neuroprosthesis. Nat. Rev. Neurosci. 25, 473–492 (2024).

2. Card, N. S. et al. An Accurate and Rapidly Calibrating Speech Neuroprosthesis. N Engl J Med 391, 609–618 (2024).

3. Metzger, S. L. et al. A high-performance neuroprosthesis for speech decoding and avatar control. Nature 620, 1037–1046 (2023).

4. Wairagkar, M. et al. An instantaneous voice-synthesis neuroprosthesis. Nature 644, 145–152 (2025).

5. Liu, Y. et al. A high-density 1,024-channel probe for brain-wide recordings in non-human primates. Nat Neurosci 27, 1620–1631 (2024).

6. Jung, T. et al. A wireless subdural-contained brain–computer interface with 65,536 electrodes and 1,024 channels. Nat Electron 8, 1272–1288 (2025).

7. Jun, J. J. et al. Fully integrated silicon probes for high-density recording of neural activity. Nature 551, 232–236 (2017).

8. Steinmetz, N. A. et al. Neuropixels 2.0: A miniaturized high-density probe for stable, long-term brain recordings. Science 372, eabf4588 (2021).

9. Zhao, Z. et al. Ultraflexible electrode arrays for months-long high-density electrophysiological mapping of thousands of neurons in rodents. Nat. Biomed. Eng 7, 520–532 (2022).

10. Trautmann, E. M. et al. Large-scale high-density brain-wide neural recording in nonhuman primates. Preprint at 10.1101/2023.02.01.526664 (2023).

11. Qian, Y. et al. Real-time decoding of full-spectrum Chinese using brain-computer interface. ScieNce AdvANceS (2025).

12. Schneider, S., Lee, J. H. & Mathis, M. W. CEBRA : Learnable latent embeddings for joint behavioural and neural analysis. Nature 617, 360–368 (2023).

13. Liu, Z. et al. A memristor-based adaptive neuromorphic decoder for brain–computer interfaces. Nat Electron 8, 362–372 (2025).

14. Shih, J. J., Krusienski, D. J. & Wolpaw, J. R. Brain-Computer Interfaces in Medicine. Mayo Clinic Proceedings 87, 268–279 (2012).

15. Wolpaw, J. R., Birbaumer, N., McFarland, D. J., Pfurtscheller, G. & Vaughan, T. M. Brain– computer interfaces for communication and control.

16. Fernandez-de-Cossio-Diaz, J. et al. Designing molecular RNA switches with Restricted Boltzmann machines. Nat Commun 16, 11223 (2025).

17. Romero, S. V. et al. Protein folding with an all-to-all trapped-ion quantum computer. Preprint at 10.48550/arXiv.2506.07866 (2025).

18. Pal, S., Bhattacharya, M., Lee, S.-S. & Chakraborty, C. Quantum Computing in the Next-Generation Computational Biology Landscape: From Protein Folding to Molecular Dynamics. Mol Biotechnol 66, 163–178 (2024).

19. Robert, A., Barkoutsos, P. Kl., Woerner, S. & Tavernelli, I. Resource-efficient quantum algorithm for protein folding. npj Quantum Inf 7, 38 (2021).

20. Inagaki, T. et al. A coherent Ising machine for 2000-node optimization problems. Science 354, 603–606 (2016).

21. McMahon, P. L. et al. A fully programmable 100-spin coherent Ising machine with all-to-all connections. Science 354, 614–617 (2016).

22. Wei, H. et al. A versatile coherent Ising computing platform. Light Sci Appl 15, 74 (2026).

23. Al-Kayed, N. et al. Programmable 200 GOPS Hopfield-inspired photonic Ising machine. Nature 648, 576–584 (2025).

24. Laydevant, J., Marković, D. & Grollier, J. Training an Ising machine with equilibrium propagation. Nat Commun 15, 3671 (2024).

25. Ackley, D. H., Hinton, G. E. & Sejnowski, T. J. A learning algorithm for boltzmann machines. Cognitive Science 9, 147–169 (1985).

26. Hinton, G. E. Training Products of Experts by Minimizing Contrastive Divergence. Neural Computation 14, 1771–1800 (2002).

27. Zhu, H. et al. Temporal coding carries more stable cortical visual representations than firing rate over time. Nat Commun 16, 7162 (2025).

28. Chowdhury, R. H., Glaser, J. I. & Miller, L. E. Area 2 of primary somatosensory cortex encodes kinematics of the whole arm. eLife 9, e48198 (2020).

29. Pei, F. et al. Neural Latents Benchmark ‘21: Evaluating latent variable models of neural population activity.

30. Churchland, M. M. et al. Neural population dynamics during reaching. Nature 487, 51–56 (2012).

31. Osindero, S. & Hinton, G. E. Modeling image patches with a directed hierarchy of Markov random fields. in Advances in Neural Information Processing Systems vol. 20 (Curran Associates, Inc., 2007).

32. Song, Y., Zheng, Q., Liu, B. & Gao, X. EEG Conformer: Convolutional Transformer for EEG Decoding and Visualization. IEEE TRANSACTIONS ON NEURAL SYSTEMS AND REHABILITATION ENGINEERING 31, 710–719 (2023).

33. Chen, C.-S., Chen, S. Y.-C., Tsai, A. H.-W. & Wei, C.-S. QEEGNet: Quantum Machine Learning for Enhanced Electroencephalography Encoding. Preprint at 10.48550/arXiv.2407.19214 (2025).

34. Chen, C.-S., Chen, S. Y.-C. & Tseng, H.-H. Exploring the Potential of QEEGNet for Cross-task and Cross-dataset Electroencephalography Encoding with Quantum Machine Learning. J Sign Process Syst 98, 5 (2026).

35. Chung, J. E. et al. A Fully Automated Approach to Spike Sorting. Neuron 95, 1381-1394.e6 (2017).

