## Supplementary Fig. S1-S5 for "Spikes meet Spins: Quantum-Native Neural Decoding for Ultra-Low-Latency Brain–Computer Interfaces"

**The PDF file includes:**

Figs. S1 to S5

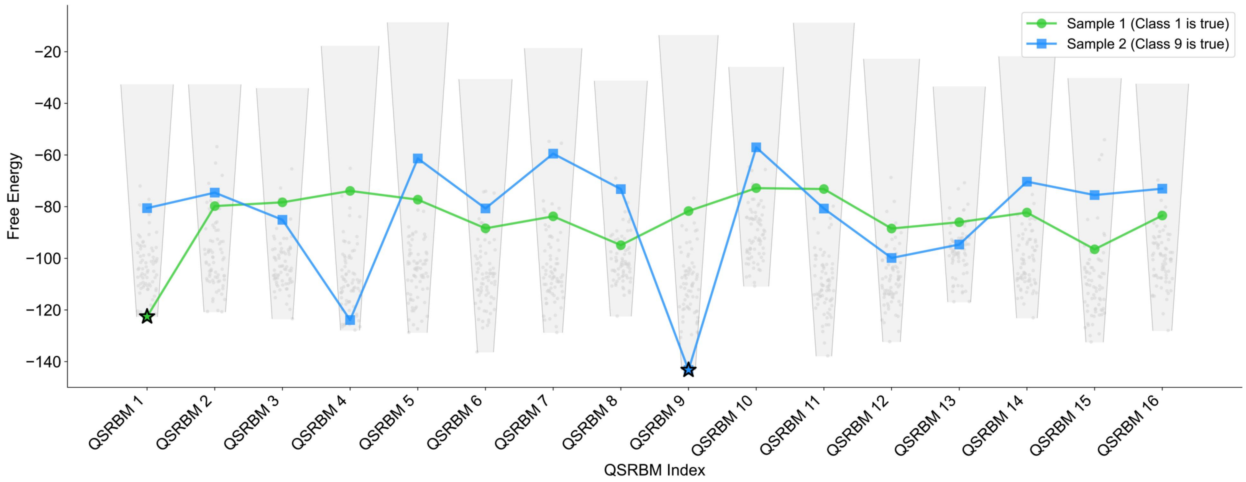


**Fig. S1. Energy funnel dynamics.**

Visualization of energy landscapes for 16-class QSRBM decoding, showing distinct energy funnels for true target classes (Sample 1: Class 1; Sample 2: Class 9). The pronounced energy gap between the true class and competing classes effectively traps non-target patterns in higher-energy states during the physical relaxation process of the photonic Ising machine, ensuring robust target classification.

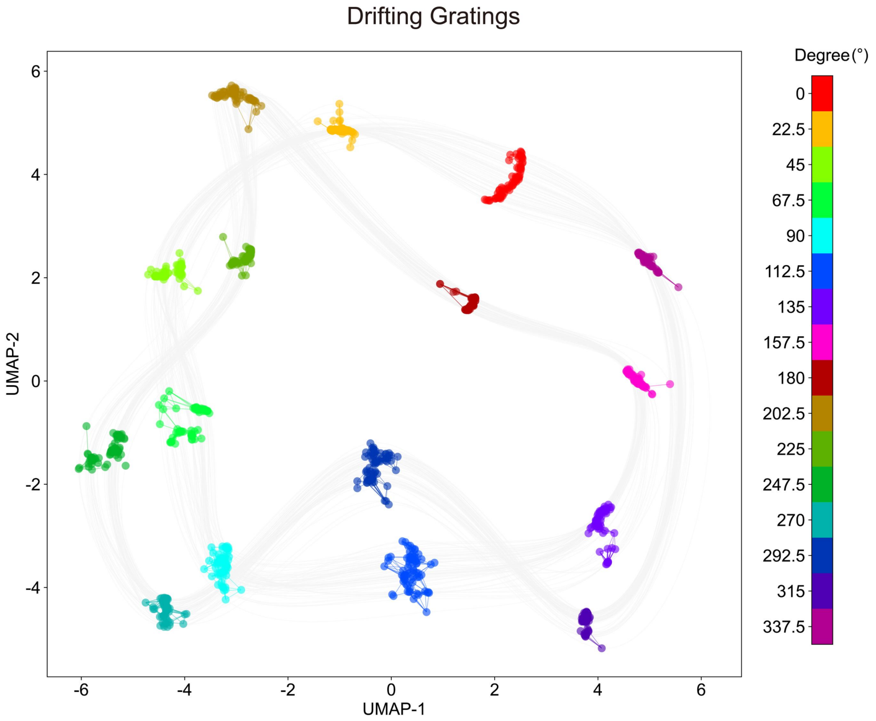


**Fig. S2. Latent energy-vector manifold for drifting grating direction decoding.**
Uniform Manifold Approximation and Projection (UMAP) visualization of energy vectors derived from the CIM for a 16-class drifting grating direction decoding task. The physically learned energy landscape generates a structured low-dimensional manifold, where energy vectors cluster regularly according to grating motion directions: not only do clusters of adjacent motion directions locate closely to each other, but those of gratings with the same orientation yet opposite motion directions also cluster in close proximity, presenting a continuous spatial geometric feature overall. Each motion direction forms a well-defined, distinctly separated cluster, enabling accurate discrimination of drifting grating stimuli.

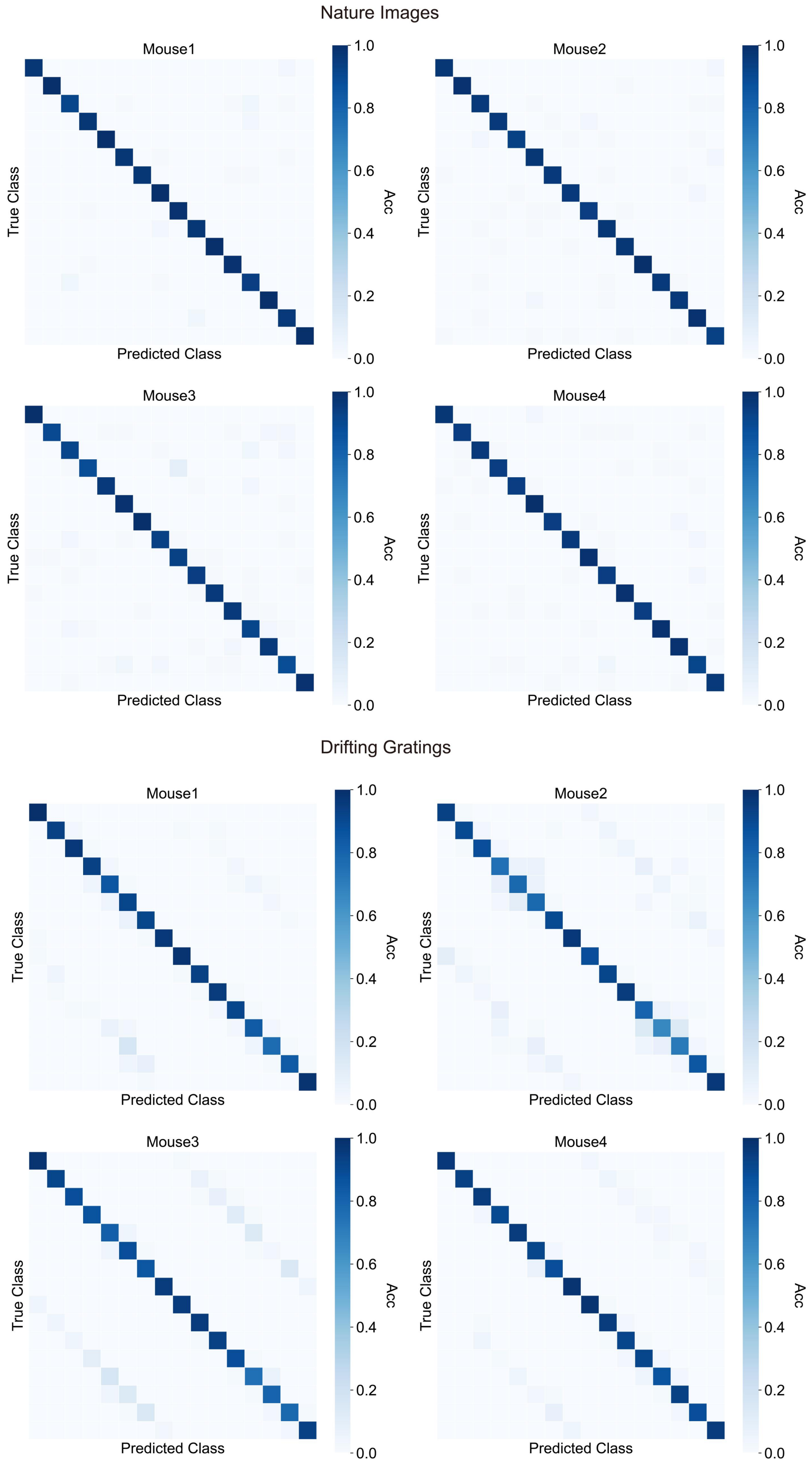


**Fig. S3. Confusion matrices for visual stimulus decoding.**

Error analysis for mouse V1 datasets (natural images and drifting gratings), showing that residual errors are largely confined to biologically similar stimuli or opposite motion directions.

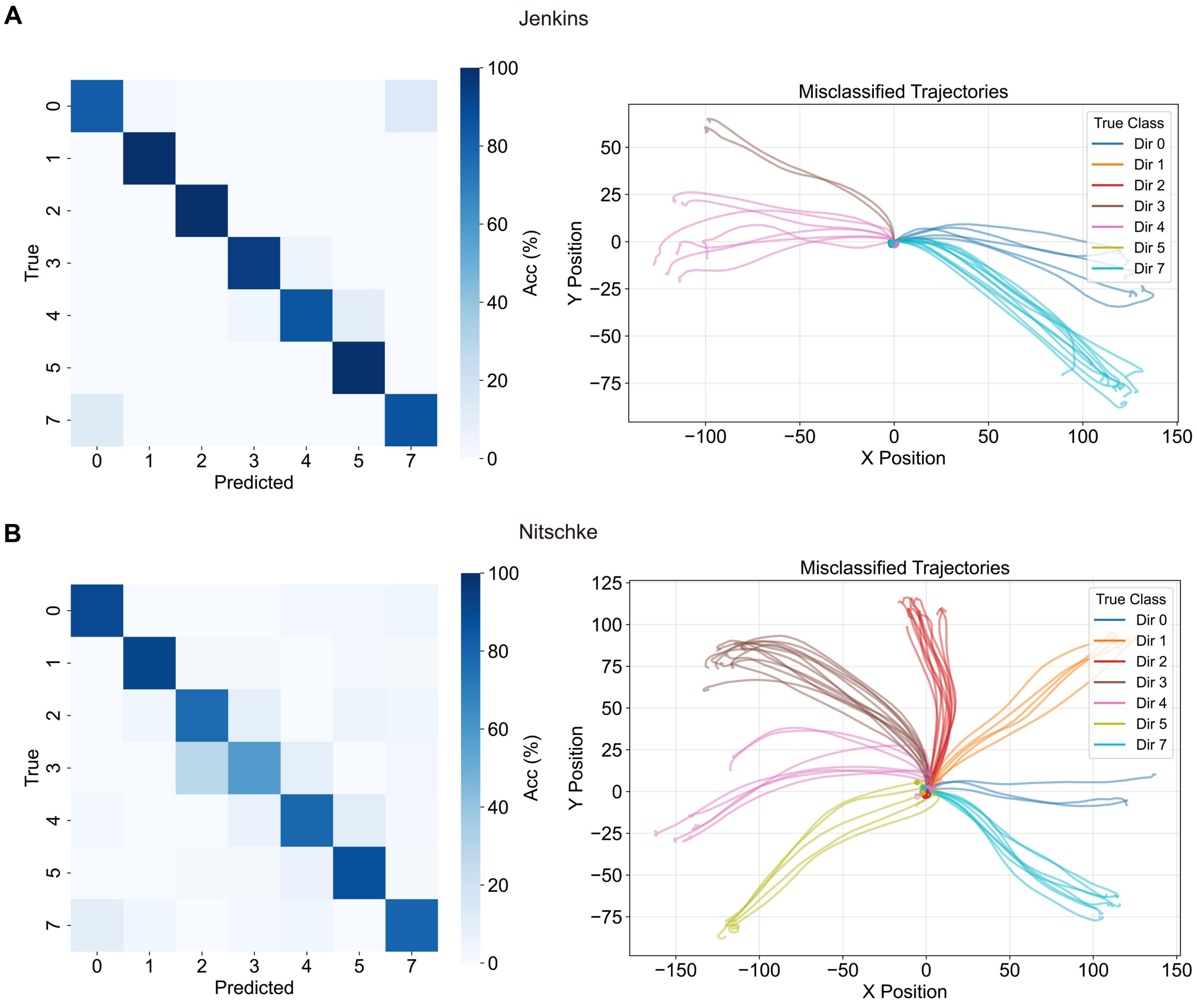


**Fig. S4. Confusion matrices and trajectory analysis for motor decoding.**

**(A, B)** Confusion matrices for cynomolgus monkeys datasets (Jenkins and Nitschke), where misclassifications primarily involve adjacent reach directions, reflecting the preservation of the directional manifold in the energy landscape.

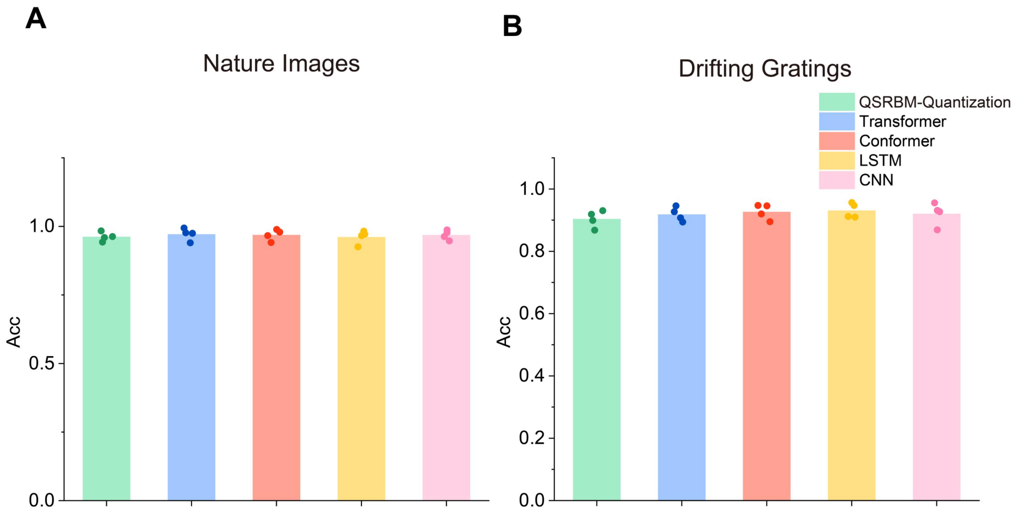


**Fig. S5. Impact of quantization on decoding performance.**

**(A, B)** Control experiment comparing the QSRBM (4-bit data) against classical models operating on raw, uncompressed spike trains, illustrating that the quantum-native approach maintains high fidelity even under extreme data compression. Data were collected from n=4 mice. Individual dots represent the mean decoding accuracy for each recording session, and bars indicate the average of these session means.
